# Constructing networks for comparison of collagen types

**DOI:** 10.1101/2023.08.25.554753

**Authors:** Valentin Wesp, Lukas Scholz, Janine M. Ziermann-Canabarro, Stefan Schuster, Heiko Stark

## Abstract

Collagens are structural proteins that are predominantly found in the extracellular matrix of multicellular animals, where they are mainly responsible for the stability and structural integrity of various tissues. All collagens contain polypeptide strands (ɑ-chains). There are several types of collagens, some of which differ significantly in form, function, and tissue specificity. Because of their importance in clinical research, they are grouped into subdivisions, the so-called collagen families, and their sequences are often analysed. However, problems arise with highly homologous sequence segments. To increase the accuracy of collagen classification and prediction of their functions, the structure of these collagens and their expression in different tissues could result in a better focus on sequence segments of interest. Here, we analyse collagen families with different levels of conservation. As a result, clusters with high interconnectivity can be found, such as the fibrillar collagens, the COL4 network-forming collagens, and the COL9 FACITs. Furthermore, a large cluster between network-forming, FACIT, and COL28a1 ɑ-chains is formed with COL6a3 as a major hub node. The formation of clusters also signifies, why it is important to always analyse the ɑ-chains and why structural changes can have a wide range of effects on the body.

## Introduction

### Collagen

Collagens are structural proteins of the extracellular matrix (ECM) in multicellular animals and, with a mass fraction of ∼30 %, constitute an essential part of the ECM’s structure and maintenance (Huxley-Jones et al., 2007; LeBleu et al., 2007; Patino et al., 2002; Salamito et al., 2021). Collagens are found in skin, connective tissue, vascular walls, bone, and cartilage (Ahmed et al., 2018; Bornstein & Sage, 1980; Kucharz, 1992; Miller & Gay, 1982; Zhao et al., 2021). Moreover, the expression of collagens is essential for the formation of the basic structure of tissues during ontogenesis (Van Der Rest & Garrone, 1991). For example, a somite is a block of condensed mesoderm formed bilaterally along the central axis in vertebrate embryos (Tajbakhsh & Spӧrle, 1998). Collagens are not only essential for the formation of these blocks but also for the differentiation of the somitic subunits, i.e., dermatome, myotome, and sclerotome (Duband & Thiery, 1987; Leivo et al., 1980). The fusion of myoblasts into multinucleated myofibers during skeletal and cardiac muscle development is another essential function of one of the collagens (Gonçalves et al., 2019).

Many types of collagens vary widely in their form, function, and tissue specificity. Currently, 28 different types of collagens are known, which can be divided into six different families (Table 1; Gordon & Hahn, 2010; Mayne & Brewton, 1993; Mayne & Burgeson, 1987; Vuorio & De Crombrugghe, 1990). A classification is based on characteristic properties of the collagen such as its structure, interaction, or site of expression (Ricard-Blum, 2010). Due to the variability of characters, various classifications of collagen exist, with none of them being indisputable (Exposito et al., 2010; Franzke et al., 2003; Knupp & Squire, 2003; Ricard-Blum et al., 2005). Thus, Table 1 is only one way of classifying collagens.

**Table 1:** Classification of the 28 collagens into six different groups based on Gordon & Hahn (2010). Collagens are colour-coded according to their group, as also used further in this article. FACITs = Fibril Associated Collagens with Interrupted Triple helices.

| Group | Member |
| --- | --- |
| Fibrillar collagens | I, II, III, V, XI, XXIV, XXVII |
| Associated with fibrils (FACITs) | IX, XII, XIV, XVI, XIX, XX, XXI, XXII |
| Network-forming collagens | IV, VI, VII, VIII, X |
| Transmembrane collagens | XIII, XVII, XXIII, XXV |
| Endostatin precursor collagens | XV, XVIII |
| Other collagens | XXVI, XXVIII |

Within the 28 collagens, 44 ɑ-chains can be distinguished. In consideration of the collagen groups in Table 1, these ɑ-chains are characterised by specific processing units that are separated by a repetitive sequence ( (Gly-X-Y)_n_ , Brodsky & Persikov, 2005), labelled in Uniprot as “Chain”. This chain is predominantly involved in structure formation. The first unit, which appears in all ɑ-chains except for the transmembrane ones, is the addition of a signal sequence that is important for the translocation of the ɑ-chain to the endoplasmic reticulum that is afterward removed (Connizzo et al., 2013; McAlinden et al., 2003). In nine fibrillar and two network-forming ɑ-chains (COL1a1, COL1a2, COL2a1, COL3a1, COL4a1, COL4a2, COL5a1, COL5a2, COL11a1, COL11a2, COL27a1), a propeptide unit is located next to the signal unit and/or at the end of all fibrillar ɑ-chains (The Uniprot Consortium, 2023). These propeptides are essential for assembling collagen trimers and are cleaved in fibrillar collagens after helix formation (Exposito et al., 2010). In addition to this, several residues are modified by post translational modifications (PTM, e.g., hydroxylation of proline). Most of the collagens are homotrimers, i.e., they are built from three identical ɑ-chains (Francomano, 1995; Brinckmann, 2005; Ricard-Blum, 2010; Gelse et al., 2003, Hulmes, 2008; Linsenmayer, 1991). However, some heterotrimers consist of two identical ɑ-chains and one different ɑ-chain. The resulting homo- and heterotrimers aggregate with collagen microfibrils in the environment, which are further organised as fibrils (Birk & Bruckner, 2005; Fratzl, 2008; Goh et al., 2014). These fibrils are further packed into fibres, which are bundled into fascicles (Heino, 2007; Nimni & Harkness, 2018).

Another characterisation of collagens is based on the abundance of their collagen types and mRNA expression throughout the human body (Kim et al., 2014; Uhlén et al., 2015). The mRNA localization of individual collagens is often tissue-specific (Oh et al., 1993; Kim et al., 2014).

The understanding of the sequence and structure of collagens is crucial because of hereditary diseases based on mutations in collagen genes (see, e.g., Arseni et al., 2018), such as osteogenesis imperfecta, Ehlers-Danlos syndrome, and Stickler syndrome.

### AlphaFold

AlphaFold DB is a protein structure database (DB) and prediction tool created by DeepMind and the European Molecular Biology Laboratory’s European Bioinformatics Institute (EMBL-EBI). It uses a deep neural network to predict spatial structures for proteins, similar to a graph interference problem (Jumper et al., 2021; Varadi et al., 2022). It maximises the number of interactions according to a graph-theoretical model considering the similarity across structures. The advantage of AlphaFold is that it can, in theory, be applied to all known amino acid sequences.

However, this requires prior measurements of 3D structures to validate these sequences, which is determined by a confidence value. This value indicates for each position in the amino acid sequences how well the prediction matches the measurement. AlphaFold categorises these confidence values (CV) into four groups concerning their accuracy. These groups are ‘high accuracy’ (CV>90), ‘generally good backbone prediction’ (70<CV<90), ‘low accuracy’ (50<CV<70), and ‘ribbon-like appearance’ (CV<50). AlphaFold suggests that the last group should not be interpreted.

Screening of the collagen sequences revealed that the proportions of repetitive regions were assigned a very low confidence value. These repetitive regions are found mainly in the inside of the sequences, as they are instrumental in the structure of the collagen (Brown & Timpl, 1995; Beck & Brodsky, 1998). However, they can complicate analysis due to their repetitions, as they occur more frequently than the functional regions of the protein. This is in contrast to boundary regions, as they return a high-confidence value in most collagens, which indicates known knowledge about the 3D structure of other proteins.

### Aim of this study

Considering the importance of collagen for ontogenesis and structure formation in the body (Marro et al., 2016; Mereness & Mariani, 2021) and the multitude of collagen-related human diseases (Arseni et al., 2018, Kuivaniemi et al. 1991), it is surprising that there are only a few complete studies regarding the possible alignment of collagens to each other (Nassa et al., 2012). In this study, the 44 α-chains of all 28 available human collagens from all six families are analysed by bioinformatics techniques.

We aim to resolve the following questions by aligning these short high-confidence regions based on their hydrophobicity. (1) Is it possible to construct informative networks between different α-chains and their collagens using high-confidence regions? (2) Are there any comparable features between the resulting networks and the literature?

## Material & Methods

### Sequence preparation

The amino acid (AA) sequence and their confidence values of human collagen variants is retrieved from the AlphaFold DB based on AlphaFold with the version from the “14th Critical Assessment of Protein Structure Prediction” (Fig. 1; Jumper et al., 2021; Varadi et al., 2022, v2.0). In AlphaFold, over 200 million protein structure predictions are stored and freely available.

**Figure 1:**
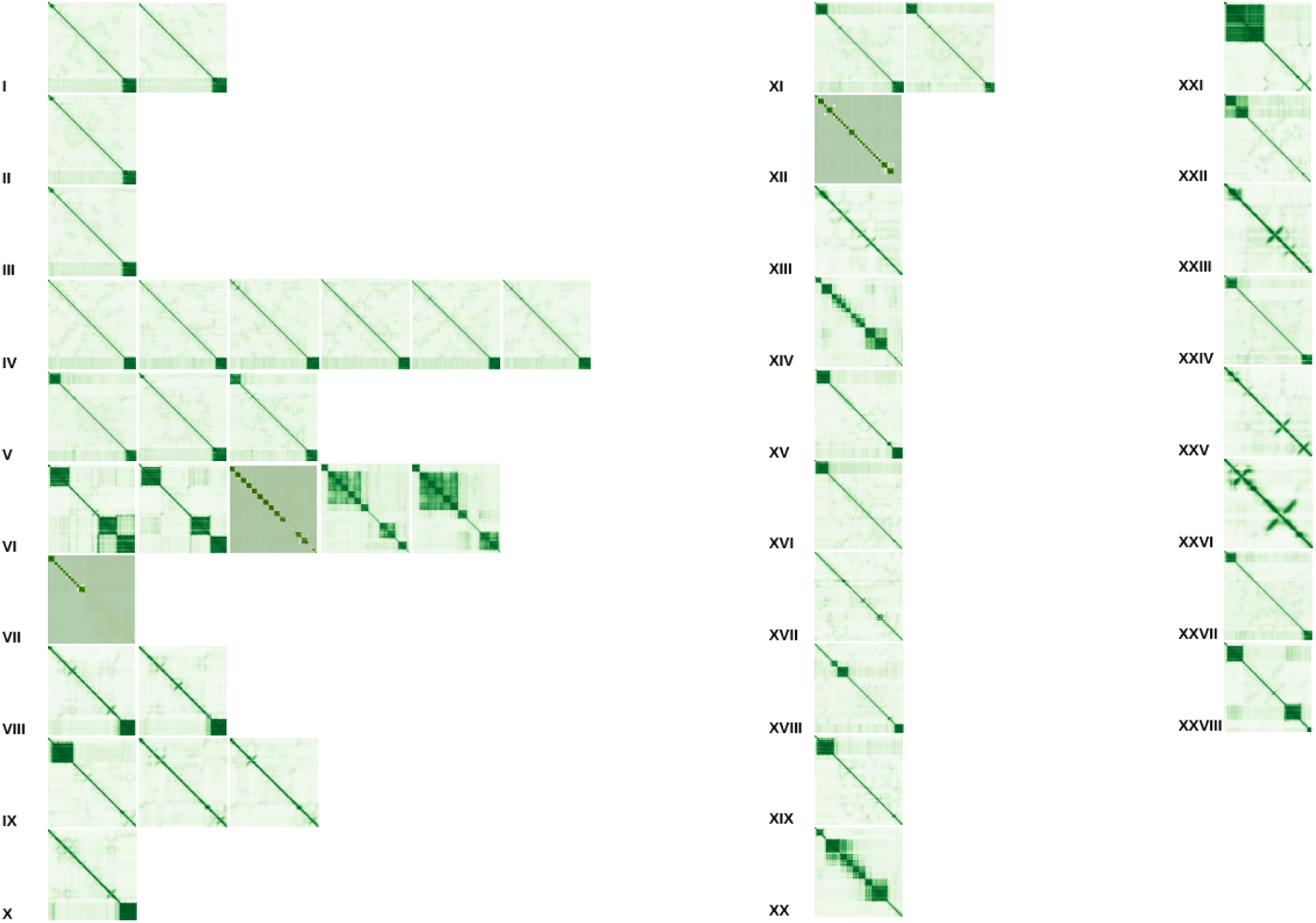
Overview of the predicted aligned error (PAE) of the collagen α-chains of collagens I-XXVIII adapted from AlphaFold (Jumper et al., 2021; Varadi et al., 2022). The colour indicates the expected position error when the predicted and true structures are aligned. The darker the colour of green, the lower the alignment error. The dark-shaded boxes were predicted with a local ColabAlphaFold instance.

The ɑ-chains are selected based on their curation state. If this is not available, the reference proteome state is favoured. The sequences and their position confidence values of the collagen α-chains are downloaded as PDB files. An exception is made for Col6a3, COL7a1, and Col12a1 for which longer and reviewed sequences are available in the UniProt database (The Uniprot Consortium, 2023). For those α-chains, we download the sequences in FASTA format and predict their confidence values with a local ColabAlphaFold instance (v.1.5.2, https://github.com/YoshitakaMo/localcolabfold). Additionally, we retrieve the PTMs for each ɑ-chain. The IDs (identification numbers) and PTM positions of the chosen ɑ-chains can be found in Supplement 1.

Since the PDB format is atypical for most sequence analysis tools, it is reformatted into a FASTA format based on the confidence values of each position. To process the following steps, several custom python (v3.10.12) scripts are implemented (Supp. 7). First, all amino acids are transferred from the 3-letter code to the 1-letter code based on the IUPAC nomenclature. Then, high-confidence regions (based on their average confidence value) from each α-chain are extracted (Fig. 2). This is done separately for different subsequence lengths (from ten AA to 500 AA in steps of ten) and confidence thresholds (from zero to 95 in steps of five). To ensure that parts of each region do not appear twice or more often, after the extraction of each subsequence the confidence values of the particular region of the α-chain are set to -1×10⁹. Furthermore, for each subsequence, the reversed complement is computed (marked with “rev” in the sequence name). This is done for three conditions based on the PTM positions of each ɑ-chain: WHOLE SEQUENCE (entire ɑ-chain sequence), WITHOUT SIGNAL (the signal sequence is removed from the ɑ-chain sequence), and ONLY CHAIN (the signal and propeptide sequences are removed from the ɑ-chain sequence).

**Figure 2:**
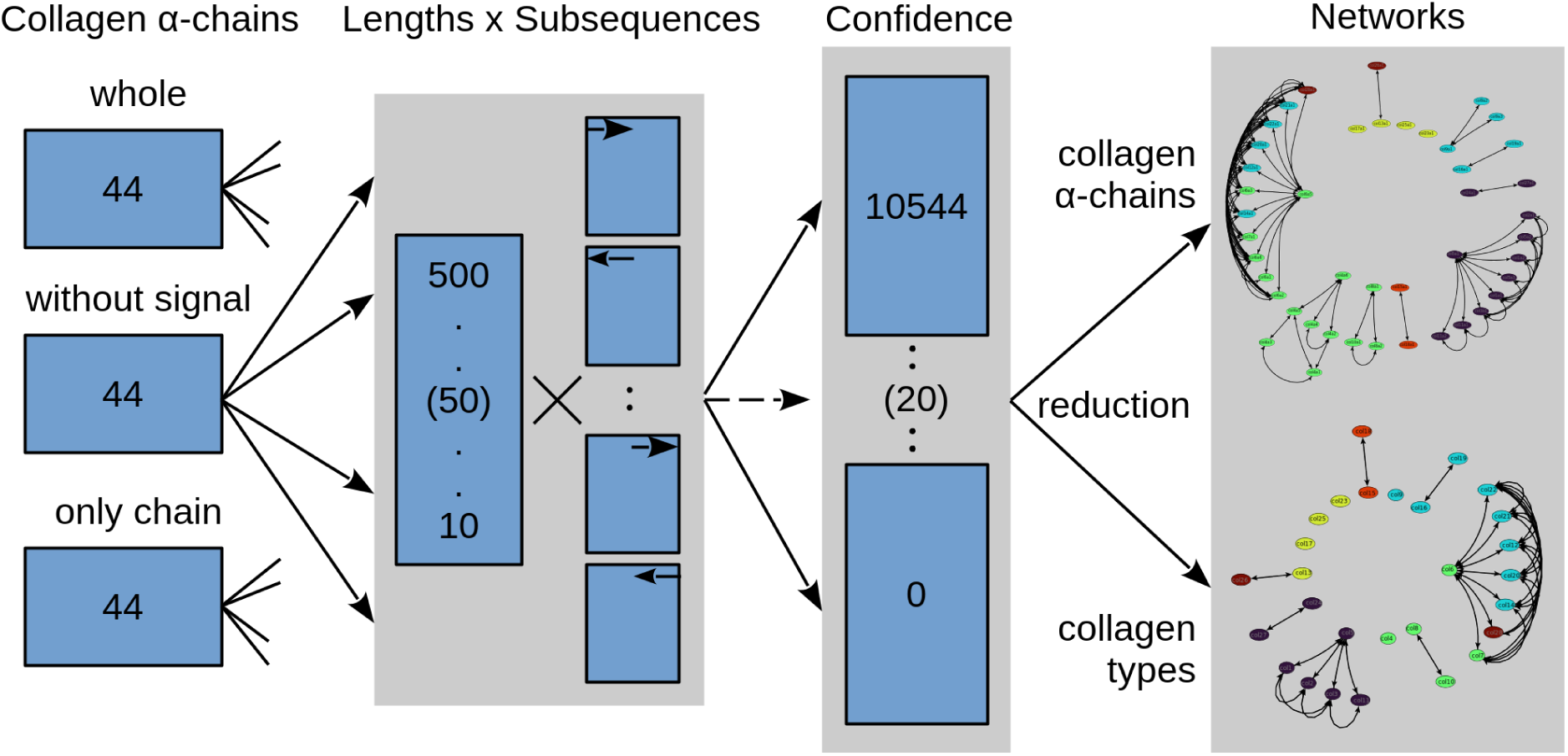
Flowchart of data processing for the 44 collagen α-chains. From left to right: we have collected 44 collagen ɑ-chains based on three conditions: WHOLE SEQUENCE (entire ɑ-chain sequence), WITHOUT SIGNAL (the signal sequence is removed from the ɑ-chain sequence), and ONLY CHAIN (the signal and propeptide sequences are removed from the ɑ-chain sequence). From these, subsequences are extracted based on lengths (between ten and 500 in steps of ten) and confidence values (between zero and 95 in steps of five). Afterwards, for each set, all sequences were globally aligned to each other and matrices were constructed based on the calculated similarities. In the end, all similarity matrices were visualised in the form of networks. Each node was coloured based on the colours in Table 1.

### Construction of similarity matrices

To create the distance similarity matrices on which we base our networks, each subsequence in a set is globally aligned to all other subsequences based on their hydrophobicity. The substitution matrix is based on the Eisenberg consensus scale, which combines several hydrophobicity (e.g., 0.48 kcal/mol for glycine or 0.12 kcal/mol for proline) features into one score (Supp. 2). The difference in hydrophobicity for each amino acid pair is taken as the substitution rate and normalised between zero and 20. The rate values are rounded to the next integer. Gap open cost is set to ten and the gap extend cost to 0.5 (EMBOSS default). Additionally, to compare each set with the others, the calculated similarities are also normalised to the length of the aligned subsequences (resulting in a maximum score of 20 if two identical sequences are compared).

### Compression of similarity matrices for further analysis

Afterward, data compressions on the similarity matrices are performed to better analyse our results. The subsequences are assigned to one of two categories, one containing all α-chains and one containing only the collagen types. For the α-chains (44), the maximum similarity between two α-chains over all their subsequence comparisons is selected. For the collagen types (28), the maximum similarity between the subsequences of all α-chains of the compared collagen types is selected. In a further step, lists are generated from these two compressed datasets listing the number of pairings between each α-chain subset and each collagen subset. For these similarities, score thresholds are set between 12 and 20 in steps of 0.25. This is done for each subsequence length and confidence value.

For each reduced similarity matrix, a network (in GML format) is built with the ɑ-chains/collagens as nodes. The similarity scores are used as edges between each α-chain and based on the score thresholds using a custom PASCAL program (v1.1.24). The networks are visualised with the general-purpose diagramming program yEd (v3.23.1).

### Generation of literature matrix for validation

To extrapolate the connection between the different α-chains, we construct a reference matrix between all α-chains based on reference databases. The idea behind this is that even if only neighbour links are considered, a larger network should still result. This is done for Google Scholar (entire text) and PubMed (only title and abstract) databases. For this purpose, a search query is performed between two collagens (Google Scholar) or α-chains (Pubmed) for all collagens and α-chains in humans. The number of hits is stored in a matrix. To avoid bias by focusing on specific collagen types in the literature, these values are logarithmized (to the base of 100) and normalised between zero and 20. As with the alignment networks, different score thresholds for the networks were used (between one and 20 in steps of one).

## Results

Here, we aligned the short high-confidence regions of collagens based on their hydrophobicity. This enabled us to investigate the potential of different collagen ɑ-chains to build informative networks. This should help reveal unknown interactions between different collagen types and make it possible to infer their spatial arrangement through simulations, i.e., within the basement membrane (e.g., basal lamina). Additionally, we compared our networks to the literature to verify their correctness.

### Identification of relevant parameters

Overall, we calculated 192,000 different networks (three sequence types [WHOLE SEQUENCE, WITHOUT SIGNAL, ONLY CHAIN], 50 subsequence lengths, 20 confidence thresholds, 32 network thresholds, and two experiments [ɑ-chains, collagens]). Therefore, to reduce the complexity of the results, we further adjusted the parameters. According to AlphaFold, a confidence threshold of at least 70 is required to have a “good backbone prediction”. In the case of the WHOLE SEQUENCE and WITHOUT SIGNAL sequence types, the longest possible subsequence length for confidence 70, for which all ɑ-chains (at least one sequence) are present, is at length 30. Additionally, we get the same parameters when we calculate an optimum over the entire set of matrices (50 subsequences and 20 confidence thresholds) between the number of subsequences and the standard deviation of the similarity scores for the WHOLE SEQUENCE and WITHOUT SIGNAL datasets. For simplicity’s sake, we take the same parameters for the ONLY CHAIN dataset.

The high-confidence sequences and results for the WHOLE SEQUENCE, WITHOUT SIGNAL and ONLY CHAIN datasets, can be found in Supplements 3, 4, and 5, respectively. Table 2 shows the three sequence types at which confidence thresholds ɑ-chains are filtered out. For the ONLY CHAIN sequence type, even at low subsequence lengths of ten and a confidence threshold of 70 most fibrillar ɑ-chains (COL1a1, COL1a2, COL2a1, COL3a1, COL11a1, COL27a1) are filtered out.

**Table 2:**
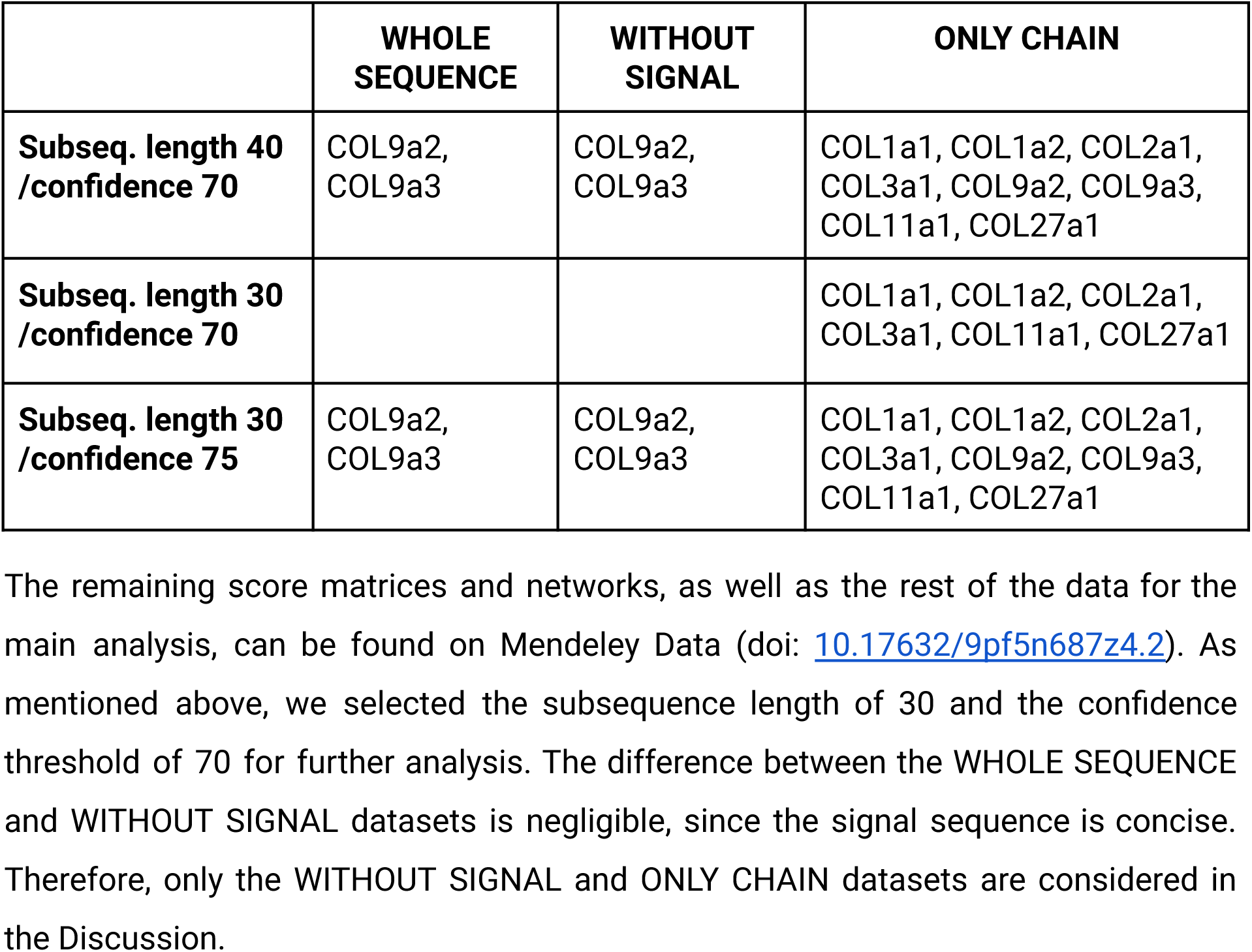
Summary of the excluded ɑ-chains based on subsequence lengths and confidence thresholds. The first row shows all ɑ-chains missing for the next highest subsequence length (40) and the third row for the next highest confidence threshold (75). The second row shows the ɑ-chains missing for our chosen parameters.

|  | **WHOLE**<br>**SEQUENCE** | **WITHOUT**<br>**SIGNAL** | **ONLY CHAIN** |
| --- | --- | --- | --- |
| **Subseq. length 40**<br>**/confidence 70** | COL9a2,<br>COL9a3 | COL9a2,<br>COL9a3 | COL1a1, COL1a2, COL2a1,<br>COL3a1, COL9a2, COL9a3,<br>COL11a1, COL27a1 |
| **Subseq. length 30**<br>**/confidence 70** |  |  | COL1a1, COL1a2, COL2a1,<br>COL3a1, COL11a1, COL27a1 |
| **Subseq. length 30**<br>**/confidence 75** | COL9a2,<br>COL9a3 | COL9a2,<br>COL9a3 | COL1a1, COL1a2, COL2a1,<br>COL3a1, COL9a2, COL9a3,<br>COL11a1, COL27a1 |

### Lists and networks

With the selected subsequence length and confidence threshold, an informative network is obtained in which the thickness of the edges indicates the similarities between ɑ-chain/collagen. For clarity, we further filtered out thin edges (comparatively low similarity) using various threshold values (from 12 to 20 in steps 0.25) to emphasise local subnetworks. From these, the first (lowest threshold) network with the most subnetworks was chosen. In all three sequence types, it was the network with the threshold of 17.5 (WHOLE SEQUENCE and WITHOUT SIGNAL: nine, ONLY CHAIN: eight). As mentioned above, for the network of the ONLY CHAIN type most fibrillar ɑ-chains are filtered out due to their repetitive chain sequences, leading to poor confidence and similarity values. This is not the case for the other two sequence types. The difference between the WHOLE SEQUENCE and WITHOUT SIGNAL datasets are negligible. Therefore, we focus on the network of the WITHOUT SIGNAL type.

As can be seen in the top panel Fig. 3, nine subnetworks are formed: (1) the fibrillar subnetwork (consisting of all fibrillar ɑ-chains, except COL24a1 and COL27a1), (2) the COL4 subnetwork (consisting of all six COL4 ɑ-chains), (3) the COL9 subnetwork (consisting of all three COL9 ɑ-chains), (4) the COL8/10 subnetwork (consisting of the two ɑ-chains of COL8 and the COL10a1 ɑ-chain), (5-8) four pair subnetworks (COL16a1 and COL19a1, COL15a1 and COL18a1, COL13a1 and COL26a1, COL24a1 and COL27a1), three unconnected transmembrane ɑ-chains (COL17a1, COL23a1, and COL25a1), and (9) the largest subnetwork consisting of the remaining ɑ-chains.

**Figure 3:**
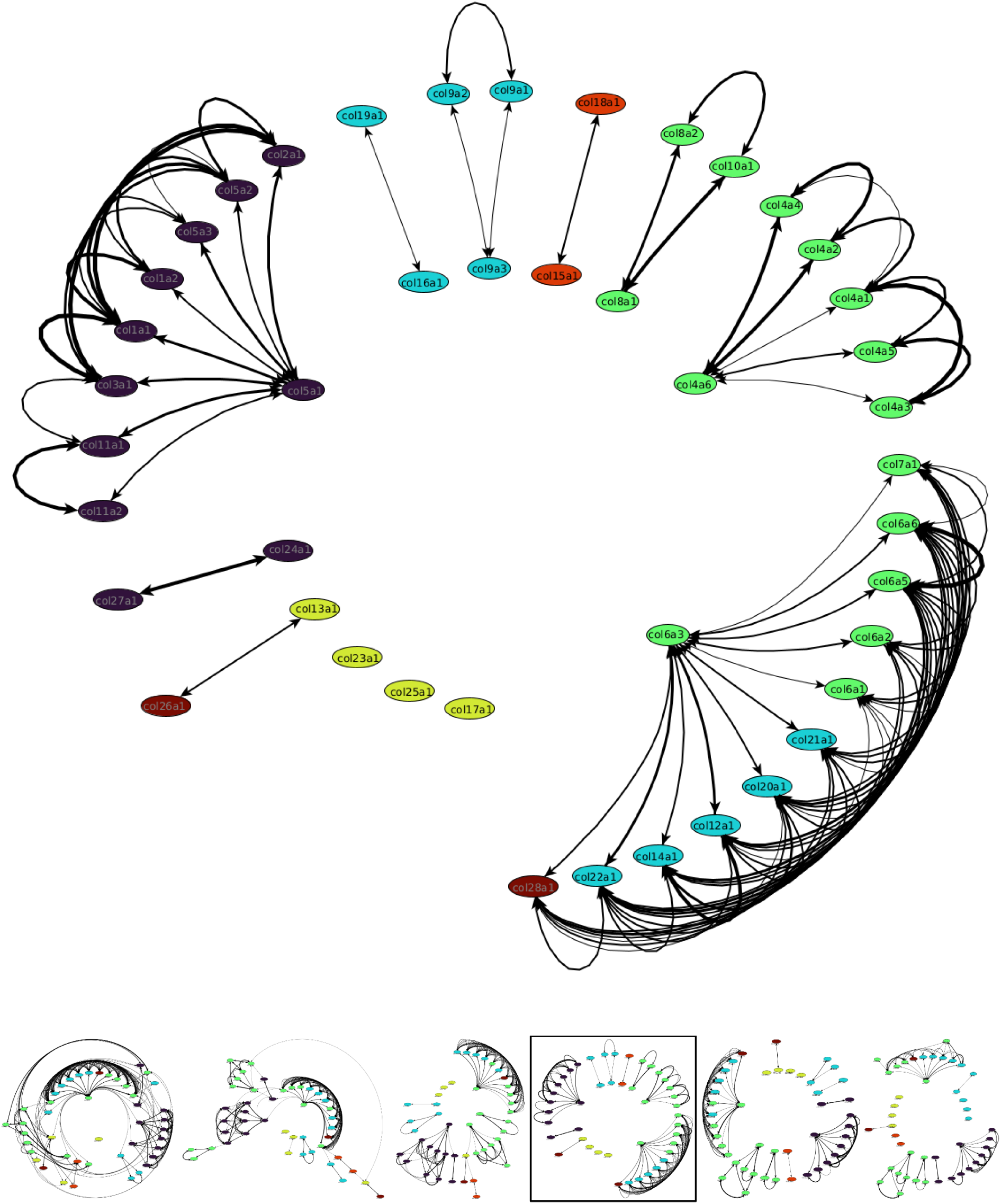
Top: Network representation of the similarity matrix (sequence type: WITHOUT SIGNAL, subsequence length: 30, confidence: 70, network threshold: 17.5) of the 44 α-chains found in humans. Bottom: From left to right, the network representation of the similarity matrix for the network thresholds 16.75, 17.0, 17.25, 17.5 (selected, same as top), 17.75, and 18.0. Colours: Purple, fibrillar collagens; cyan, associated with fibrils (FACITs); green, network-forming collagens; yellow, transmembrane collagens; orange, endostatin precursor collagens; red, other collagens.

To show the interconnection between the individual subnetworks, it is necessary to lower the threshold to 17.25. With this, the following new connections are formed (Fig. 3 bottom): The pair subnetworks are incorporated into the larger subnetworks. COL24/27 is connected to the fibrillar subnetwork through COL1a2, COL13/26 to the COL4 subnetwork through COL4a4, and COL15/18 to COL13/26 through COL13a1. At even lower thresholds (16.75, 17.0), almost all subnetworks are connected. The fibrillar network is connected to the COL4 subnetwork through COL4a3 and COL4a4, the COL8/10 subnetwork through COL8a2, and the largest subnetwork through COL6a5 and COL7a1. The subnetworks COL15/18 and COL16/19 now connect to the largest subnetwork through COL6a6. A new pair subnetwork between COL23a1 and COL25a1 is formed. Interestingly, the COL9 subnetwork is very stable.

With even lower thresholds, the networks become multidimensional and are difficult to visualise.

### Network comparison with PubMed and Google Scholar

Looking at the reference matrices (Supp. 6), the lowest values (and thus the most hits) between collagens (Google Scholar) or α-chains (PubMed) were on the diagonal, where only single collagens or α-chains were searched for. This is because single keywords always have more hits than the conjunction with other keywords. The overall lowest value was for the collagen COL1 with 0.32 (13100 hits) or the α-chain COL1a1 with 0.13 (2264 hits) indicating a high impact in the literature. The second-lowest values were for the cells around the diagonal. These cells mostly represented α-chains of the same or closely related collagens. For example, the lowest values for the six α-chains of COL4 were for each of these chains with itself. None of those values were above 0.42 and, therefore, could be found together in the literature. This could indicate that they are closely related or interact with each other. The same correlation could be seen in the networks for the alignments, where the α-chains of COL4 always formed a subnetwork with each other, but only with other α-chains at lower thresholds.

Similar results could be seen for the fibrillar collagens. In general, the values were around 0.5 or lower. However, exceptions exist due to the few literature available for specific ɑ-chains. For example, COL5a3 has values above one or even a value of ten (meaning an occurrence of zero) with COL1a2. The same can also be seen for COL24a1 and COL27a1. The lowest value was 0.72 between COL27a1 and COL1/COL5a1 while most other values were ten. At the same time, our network shows the same pattern by separating the COL24/27a1 pair from the large fibrillar network at higher thresholds. In contrast to the alignment analysis, the PubMed networks tend to form a single large cluster, even at high thresholds (Fig. 4). Further analyses were only done for PubMed, as the query limit for Google Scholar was exceeded for ɑ-chains.

**Figure 4:**
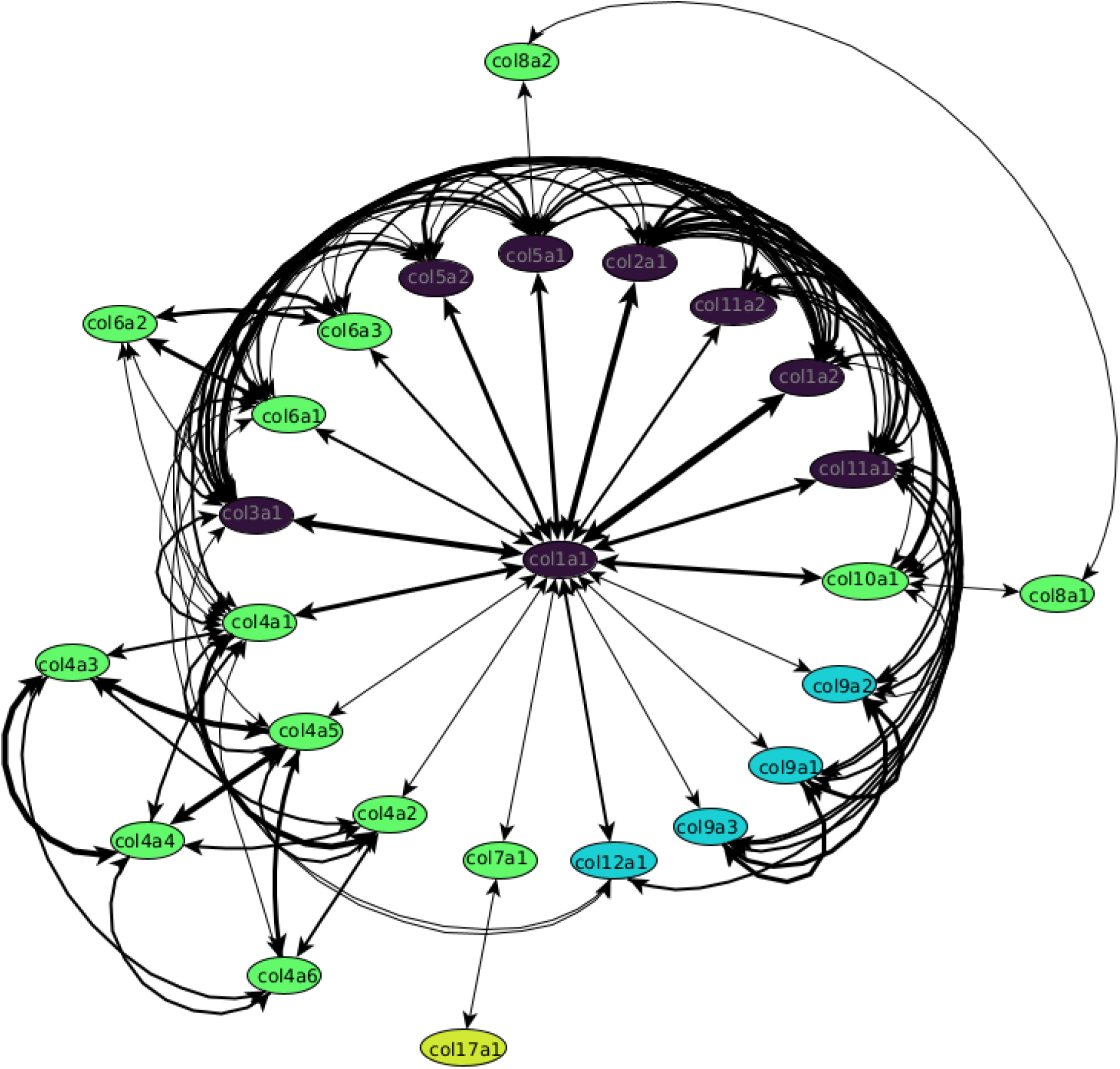
Network representation of the reference matrix of the 26/44 α-chains with significant entries found in PubMed. The other 18/44 ɑ-chains had no connections and were not shown. The colours are as follows: Purple: fibrillar collagens, cyan: associated with fibrils (FACITs), green: network-forming collagens, and yellow: transmembrane collagens.

## Discussion

We aimed to construct informative networks between the different collagen ɑ-chains based on pairwise sequence alignments of high-confidence regions and compared them with the literature. For reasons of complexity, we focus our analysis on collagens only and exclude other binding proteins. Here, we showed that this classification could be generally reconstructed again based on the sequences of the α-chains.

### Fibrillar-forming collagen clusters

It is striking that the propeptides are essential for this analysis. Without them, fibrillar collagens are filtered out due to the poor confidence values of the chain sequences. In a more detailed analysis of our networks, it can be seen that the fibrillar collagens form two distinct clusters. It turned out that COL5a1 is the major hub node in the larger cluster, which can be explained by its role as a regulator for the formation of a uniformly small corneal fibril diameter with COL1 (Mak et al., 2016). COL5a1/a2 are associated with several diseases, including Ehlers-Danlos syndrome, which is characterised by very elastic skin, weakened blood vessels, and joint hypermobility (Imamura et al., 2000; Steinmann et al., 2002; Malfait et al., 2010). COL24a1 and COL27a1 form their network separate from the other fibrillar collagens starting at a threshold of 17.5 (Fig. 3). The triple helices of these two α-chains are shorter than those of the other fibrillar collagens (Gordon & Hahn, 2010). This may be why they do not interact with other collagen ɑ-chains at high thresholds.

### Network-forming collagen clusters

The α-chains of the network-forming collagen COL4 are closely related, which indicates similar functions with few differences. As with the collagens of the fibrillar cluster, the α-chains of COL4 are also essential structure proteins in the body (Khoshnoodi et al., 2008). At a lower threshold of 17.0, a connection between COL4 and COL7 can be seen as mentioned in the literature (Roig-Rosello and Rousselle, 2020; Brittingham et al., 2006). A defect of the α-chains COL4a3/a4/a5 can lead to Alport syndrome which affects the basement membranes of the kidney, inner ear, and eye (Gregorio et al., 2023; Imafuku et al., 2018; Shulman et al., 2021; Deltas, 2022). The second cluster is formed by COL8a1, COL8a2 and COL10a2. This can be explained by their high similarity to each other, with COL8a1/a2 having one additional exon compared to COL10a1 (Gordon & Hahn, 2010). Furthermore, COL8a1 and COL10a1 are associated with age-related macular degeneration (Cascella et al., 2018) whereas mutations in COL8a2 can lead to Fuchs endothelial dystrophy, an impairment of vision (Zhang & Patel, 2015). The remaining network-forming collagens are part of the largest network, where COL6a3 is a major hub node.

### Largest cluster

The largest cluster is not as homogeneous as the previous two clusters, consisting of three different types of collagens (network-forming, FACITs, and COL28a1). All ɑ-chains are highly connected to each other. In this cluster, COL6a3 is a major hub node. Generally, COL6 forms the most connections with other collagens. At lower network thresholds (17.0 and lower), the ɑ-chains of COL6 seem to be a major hub node for the whole network (Fig. 3). This could be explained by the fact that COL6 is important for the function and stability of the cell membrane by interacting with other collagens such as COL1, COL2, COL4, and COL14 (Cescon et al., 2015; Tonelotto et al., 2019). Such connections can also be seen in our network for different thresholds, at lower thresholds for COL1/2 (17.0) and COL4 (16.75) and for COL14 at even higher thresholds of 18.0 (Supp. 4).

This network seems to contain important collagens for developmental mechanisms. One disease associated with this is limb-girdle muscular dystrophy. This disease can be caused by mutations of multiple genes encoding for proteins within the sarcolemma, cytosol, or nucleus of a myocyte (muscle cell), leading ultimately to membrane instability, a weakness of the dystrophin associated glycoprotein complex, and defects in muscle repair mechanisms (Bushby et al., 2014, Murphy & Straub, 2015). The muscular weakness is caused by destabilisation of the structural proteins that are supposed to keep the muscular cell intact during contractions; one of these proteins could be those of COL6 (Dowling et al., 2021).

### COL9 cluster

All three COL9 ɑ-chains are relatively short (the longest chain, COL9a1, is only 921 AA long) and highly similar to each other. It is known that defects in these α-chains can lead to Stickler syndrome, a disease characterised by ophthalmic, orofacial, articular, and auditory defects (Robin et al., 2000). The range of defects highly suggests that the α-chains of COL9 are expressed in the first and second pharyngeal arch derivatives, which include, for example, the maxilla, mandible, palate, and auditory ossicles (Trainor & Krumlauf, 2001, Liu et al., 2013). This could also explain why the ɑ-chains of COL9 seem to only connect to other ɑ-chains at low network thresholds (Fig. 3).

### Pair clusters

One cluster consisting of only two ɑ-chains is COL15/18. COL15a1 is structurally homologous to COL18a1 (Marneros & Olsen, 2001) which explains their distinct and stable bond. COL18a1 is mostly expressed in the brain and eye, and associated with the Knobloch syndrome, which leads to eye deformations in the development phase, called occipital encephalocele (Seppinen & Pihlajaniemi, 2011). Mouse mutants deficient in COL15a1 showed progressive degeneration in skeletal muscles and susceptibility to muscle damage (Marneros & Olsen, 2001). Other than that, no discernable feature can be extracted from the network.

For the other two pair clusters COL13/26 and COL16/19 the literature search results in no entries concerning interactions. Therefore, no information could be extracted.

### Comparison with the literature

Our results regarding the similarity matrix are also reflected in the PubMed reference matrices. In particular, local clusters can be detected. COL1a1 forms the main hub node, with the remaining fibrillar collagens having the strongest connection to each other. At the same, the COL4 and to a lesser degree the COL6, COL8/10 and COL9 clusters also emerge as subclusters in the network. However, it should be noted that this is not surprising, since ɑ-chains and collagens of the same type are usually studied together. The same can be said for COL1a1 functioning as the main hub node. COL1 is one of the most abundant structural proteins in vertebrates and, therefore, often used in conjunction with other ɑ-chains in studies (Stover and Verrelli, 2011). Lastly, it should be noted that the number of references decreases considerably at higher collagen numbers. For the ɑ-chains not shown, future studies will result in higher literature data and may remedy this problem. Overall, the resulting reference network was as expected, but could still be used to validate parts of our alignment network.

With this study, we constructed an informative network and compared it with the literature. We were also able to link this network to specific questions (i.e., regarding diseases and developmental biology).

### Outlook

Since collagens are important structural proteins present in every tissue of multicellular animals, an alteration naturally leads to serious effects on the body (Iozzo & Gubbiotti, 2018). Our analysis showed that the representation with only collagen types is not sufficient to show the linkages, because the α-chains bind differently within a collagen type. To this end, it is necessary to always examine the α-chains. Moreover, in further studies, it should be possible to verify the overall network and its subnetworks of collagens with immunohistochemistry. Additional analysis could examine the ageing of collagens (regarding chemical modifications) and its effects on networks (Fichtner et al., 2020); or also the consideration of further protein alignments between the collagens, such as adhesion proteins like fibronectins, laminins, tenascins, or glycoproteins, or in the far future for the whole human proteome.

## Supporting information

This file contains a short description of each supplement file.

This file contains the alpha-chain IDs and PTM positions.

This file contains the hydrophobicity substitution matrix.

This zip-file contains the data and figures of the WHOLE SEQUENCE dataset.

This zip-file contains the data and figures of the WITHOUT SIGNAL dataset.

This zip-file contains the data and figures of the ONLY CHAIN dataset.

This zip-file contains the data and figures of the literature reference dataset.

This zip-file contains all scripts used in this study.

## Acknowledgments

We thank all the members of the Department of Bioinformatics who were providing guidance and support during this study. Additionally, we thank A. Berndt for stimulating discussions. This study benefited from the research experience of HS gained in studies funded by the Center of Interdisciplinary Prevention of Diseases related to Professional Activities (KIP) funded by the Friedrich-Schiller-University Jena and the ’Berufsgenossenschaft Nahrungsmittel und Gastgewerbe Erfurt (Germany)’ (BGN).

## Author contributions

V.W., L.S., and H.S. conceived the study. V.W. and H.S. supervised and performed the calculations. V.W. and H.S. analysed network data. V.H. and H.S. drafted the manuscript and all figures. J.M.C.Z. contributed developmental and muscular functional aspects. S.S contributed biochemical and bioinformatics expertise. All authors contributed to the interpretation of the results and revised the manuscript.

## Mendeley repository

The datasets generated and analysed during this study along with the code and several supplementary files are available in the Mendeley repository: Wesp, Valentin; Stark, Heiko (2024), “Predicting bonds between collagens – An alignment approach”, Mendeley Data, V3, doi: 10.17632/9pf5n687z4.3

