## Supplementary material for "Constructing networks for comparison of collagen types": This zip-file contains all scripts used in this study.: CheckMultifastas.py

```
import sys
import os


def main(mfaa_files, pdb_files):
    # get all multifasta files
    multifastas = [x for x in os.listdir(mfaa_files) if x.endswith(".mfaa")]
    # get all collagen names from the PDB files
    collagens = [os.path.splitext(os.path.basename(x))[0] for x in
                 os.listdir(pdb_files) if x.endswith(".pdb")]
    # loop over all multifasta files
    for mfaa in multifastas:
        # list containing all collagens that weren't found in the multifasta
        # file
        not_found_collagens = []
        # read te multifasta file
        with open(os.path.join(mfaa_files, mfaa), "r") as reader:
            content = reader.readlines()
            # loop over all collagens
            for collagen in collagens:
                # variable checking if the collagen was found in any form
                # (left, right and reversed)
                collagen_found = False
                # loop over all lines in the multifasta file
                for line in content:
                    # check the header of each sequence if it contains the given
                    # collagen
                    # if it does, go to the next collagen and check it
                    if line.startswith(">") and collagen in line:
                        collagen_found = True
                        break

                # if the collagen wasn't found, save it
                if not collagen_found:
                    not_found_collagens.append(collagen)

        # save all collagens in a text file that weren't found
        output = os.path.join(mfaa_files, mfaa.replace(".mfaa", "_missing.txt"))
        with open(output, "w") as writer:
            writer.write("\n".join(not_found_collagens))


if __name__ == "__main__":
    # user specifies the folder with the multifasta files and the folder with
    # the PDB files
    main(sys.argv[1], sys.argv[2])
```
