## Supplementary material for "Constructing networks for comparison of collagen types": This zip-file contains all scripts used in this study.: CreateScoreMatrices.py

```
import os
import sys
from Bio import SeqIO
from Bio import Align
from Bio.Align import substitution_matrices
import pandas as pd
import natsort as ns
import multiprocessing as mp


def main(in_dir, out_dir, model, length, set_prog, set_size, lock):
    # create the output folder
    os.makedirs(out_dir, exist_ok=True)
    # initialize pairwise aligner
    aligner = Align.PairwiseAligner()
    # set aligner to global alignment
    aligner.mode = "global"
    # set gap open cost (EMBOSS Needle default)
    aligner.open_gap_score = -10
    # set gap extend cost (EMBOSS Needle default)
    aligner.extend_gap_score = -0.5
    # load the substitution matrix for the alignment
    aligner.substitution_matrix = substitution_matrices.load(model)
    # get all multifasta files and sort them after their confidence values
    mfaa_files = ns.natsorted([x for x in os.listdir(in_dir)
                               if x.endswith(".mfaa")])
    if(set_size == None):
        set_size = len(mfaa_files)

    # loop over all multifasta files
    for file in mfaa_files:
        # get the file name of the multifasta file
        filename = ".".join(file.split(".")[:-1])
        # get the confidence value of the multifasta file
        conf_val = "conf_" + file.split("_")[1]
        # create the output folder for each confidence value
        out_conf = os.path.join(out_dir, conf_val)
        os.makedirs(out_conf, exist_ok=True)
        # path to the multifasta file
        mfaa_file = os.path.join(in_dir, file)
        # load the multifasta file
        records = list(SeqIO.parse(mfaa_file, "fasta"))
        progress = 0
        if(len(records)):
            # create a dictionary between each entry in the multifasta file
            # it is basically a 2d-array for each entry with each other
            sc_matrix_dic = {rec_1.id:{rec_2.id:-1 for rec_2 in records}
                             for rec_1 in records}

            # loop over the entries
            for i in range(0, len(records)):
                for j in range(i, len(records)):
                    # calculate the alignment score between the two entries and
                    # save it, normalized to the length of the sequences
                    al_score = float(aligner.score(records[i].seq, records[j].seq))
                    sc_matrix_dic[records[i].id][records[j].id] = al_score / length
                    sc_matrix_dic[records[j].id][records[i].id] = al_score / length

                progress += 1
                print("\rlength_{} : {} -> current set: {:.2f}%".format(length,
                                   conf_val, 100*progress/len(records)), end="")

            # create the dataframe from the dictionary
            dataframe = pd.DataFrame.from_dict(sc_matrix_dic)
            # write the dataframe to the output CSV file
            dataframe.to_csv(os.path.join(out_conf, "score_matrix.csv"), sep="\t")

        with lock:
            set_prog.value += 1
            print("\r"+" "*100, end="")
            print("\rlength_{} : {} -> current set: 100% --> overall set: {:.2f}%".format(length,
                                         conf_val, 100*set_prog.value/set_size))


if __name__ == "__main__":
    mp.freeze_support()
    # set the multiprocessing manager
    manager = mp.Manager()
    # set the lock for multiple processes
    lock = manager.Lock()
    set_prog = manager.Value("i", 0)

    # the user specifies the input folder with the *.mfaa files, the output
    # folder, the substitution model ("HYDRO" for hydrophobicity) and the length
    # of each sequence
    main(sys.argv[1], sys.argv[2], sys.argv[3], int(sys.argv[4]), set_prog,
         None, lock)
```
