## Supplementary material for "Constructing networks for comparison of collagen types": This zip-file contains all scripts used in this study.: run_pipeline.py

```
import os
import sys
import ThresholdCutter as tc
import CheckMultifastas as cm
import CreateScoreMatrices as csm
import pandas as pd
import numpy as np
import multiprocessing as mp
from functools import partial


def align_process(seq_len, confs, pdbs, mods, out, model, set_prog, set_size,
                  lock):
    # path to the output folder for the given subsequence length
    len_dir = os.path.join(out, "subseq_len_{}".format(seq_len))
    # path to the output multifasta files
    mfaa_output = os.path.join(len_dir, "high_conf_regions")
    # loop over all confidence values
    for conf in confs:
        # run the ThresholdCutter.py script to get the subsequences
        tc.main(pdbs, seq_len, conf, mods, mfaa_output)

    # run the CheckMultifastas.py script to check for missing collagens
    cm.main(mfaa_output, pdbs)
    # path to the output folder with the score matrix files
    network_output = os.path.join(len_dir, "high_conf_networks")
    # run the CreateScoreMatrices.py script to calculate the scores
    # between all sequences in the mfaa files
    csm.main(mfaa_output, network_output, model, seq_len, set_prog, set_size,
             lock)


if __name__ == "__main__":
    mp.freeze_support()
    # set the multiprocessing manager
    manager = mp.Manager()
    # set the lock for multiple processes
    lock = manager.Lock()
    set_prog = manager.Value("i", 0)

    # path to the folder with the PDB files
    pdb_dir = sys.argv[1]
    # path to the output folder for the entire pipeline
    out_dir = sys.argv[2]

    # path to the collagen modification file
    # set all non avaiable cells to -1
    mod_frame = pd.read_csv(sys.argv[3], sep=",", index_col=0)
    mod_frame = mod_frame.fillna(-1)
    # save the position of modifications for the ThresholdCutter.py script
    # without_signal means that the signal sequence is removed from the
    # alpha-chain sequence if avaiable
    # only_chain means that only the chain sequence is taken from the alpha-chain
    # sequence
    # else, the entire sequence is taken
    mod_type = sys.argv[4]
    col_mods = None
    if(mod_type == "without_signal"):
        col_mods = {index:{"start":int(row["Signal"].split("-")[1]),
                           "end":row["Length"]} if row["Signal"] != -1
                           else {"start":0, "end":row["Length"]}
                    for index,row in mod_frame.iterrows()}
    elif(mod_type == "only_chain"):
        col_mods = {index:{"start":int(row["Chain"].split("-")[0])-1,
                           "end":int(row["Chain"].split("-")[1])}
                           for index,row in mod_frame.iterrows()}
    elif(mod_type == "whole_sequence"):
        col_mods = {index:{"start":0, "end":row["Length"]}
                    for index,row in mod_frame.iterrows()}
    else:
        raise Exception("Specify modification type")

    # minimum (starting point) length for each subsequence
    # maximum (ending point) length for each subsequence
    # step size for each subsequence length
    sub_seq_lengths = np.arange(int(sys.argv[5]), int(sys.argv[6])+1,
                                int(sys.argv[7]))
    np.random.shuffle(sub_seq_lengths)

    # minimum confidence value (starting point) for each dataset
    # maximum confidence value (ending point) for each dataset
    # step size for each confidence value
    conf_thresholds = np.arange(int(sys.argv[8]), int(sys.argv[9])+1,
                                int(sys.argv[10]))
    np.random.shuffle(conf_thresholds)

    # size of the entire set to the analyzed
    set_size = len(sub_seq_lengths)*len(conf_thresholds)

    # the substitution model to be used
    model = "HYDRO"

    # set number of threads
    threads = int(sys.argv[11])

    # start the multicore process for a given number of threads
    with mp.Pool(processes=threads) as pool:
        # run the process for the given parameters
        pool_map = partial(align_process, confs=conf_thresholds, pdbs=pdb_dir,
                           mods=col_mods, out=out_dir, model=model,
                           set_prog=set_prog, set_size=set_size, lock=lock)
        process = pool.map_async(pool_map, sub_seq_lengths)
        pool.close()
        #print(process.get())
        pool.join()

    print()
```
