## Supplementary material for "Constructing networks for comparison of collagen types": This zip-file contains all scripts used in this study.: ThresholdCutter.py

```
import sys
import statistics as st
import os
import numpy as np
import pandas as pd
import math
import natsort as ns


# dict for converting the 3-letter AA code into the 1-letter AA code
aa_code = {
    "MET": "M",
    "LYS": "K",
    "LEU": "L",
    "ARG": "R",
    "GLY": "G",
    "VAL": "V",
    "SER": "S",
    "ALA": "A",
    "PHE": "F",
    "TRP": "W",
    "GLN": "Q",
    "PRO": "P",
    "GLU": "E",
    "CYS": "C",
    "TYR": "Y",
    "ASP": "D",
    "ILE": "I",
    "HIS": "H",
    "THR": "T",
    "ASN": "N",
    "SEC": "U",
    "PYL": "O",
}


# method for getting all averages in an array in a window of a given size
def moving_average(arr, window):
    return np.convolve(arr, np.ones(window), "valid") / window


def main(folder, window, threshold, mods, output):
    # if the output folder does not exist, create it
    os.makedirs(output, exist_ok=True)
    # get all PDB files and sort them
    pdb_files = ns.natsorted([x for x in os.listdir(folder)
                              if x.endswith(".pdb")])
    all_sequences = []
    # loop over all PDB files in the folder
    for file in pdb_files:
        # open the PDB file
        with open(os.path.join(folder, file), "r") as reader:
            # variable for checking if the confidence values are found
            found_model = False
            # the current ID of the AA
            id = None
            # list containing the AAs of the PDB sequence
            seq = []
            # list containing the confidences of the PDB sequence
            conf = []
            # loop over all lines of the PDB file
            for line in reader:
                # if the "MODEL" tag is reached, the next lines will have
                # the confidences
                # if multiple models exists, only the first one is taken
                if line.startswith("MODEL"):
                    found_model = True
                # if the "ENDMDL" tag is reached, we are done with the
                # current model and can close the file
                elif line.startswith("ENDMDL"):
                    break

                # if we reached our model and a line starts with "ATOM", we
                # can extract the confidence values for each amino acid
                # position
                elif found_model and line.startswith("ATOM"):
                    line = line.strip()
                    # if the current ID doesn't match the next ID, we
                    # reached a new amino acid and can save it and its
                    # confidence
                    if id != line[21:26]:
                        # the ID of each position
                        id = line[21:26]
                        # save the current amino acid
                        seq.append(aa_code[line[17:20]])
                        # save the confidence of the current amino acid
                        conf.append(float(line[61:66]))

            conf = np.asarray(conf)
            # file name of the PDB file
            filename = os.path.splitext(os.path.basename(file))[0]
            seq = seq[mods[filename]["start"]:mods[filename]["end"]]
            conf = conf[mods[filename]["start"]:mods[filename]["end"]]
            # if the number of confidences (and therefore the sequence
            # length) is greater than the window size
            if(len(conf) >= window):
                # maximum number of possible subsequences
                max_seq_num = math.ceil(len(seq) / window)
                # get a number of non-overlapping subsequences with the
                # highest average confidence for a given window size
                for j in range(max_seq_num):
                    # get the averages over all confidences for the window
                    conf_averages = moving_average(conf, window)
                    # get the start index of the maximum confidence window
                    start = np.argmax(conf_averages)
                    # get the average value of the maximum confidence window
                    max_conf = np.max(conf_averages)
                    # check if the maximum average confidence is higher than
                    # the value
                    if(max_conf >= threshold):
                        # set all confidence values in the window for the
                        # maximum average confidence value to -10^9, so that
                        # in the next loop this region will be ignored for
                        # the average confidences
                        np.put(conf, np.arange(start, start+window, 1),
                               -10**9)
                        # get the subsequence for this region
                        high_conf_seq = "".join(seq[start:start+window])
                        # set the file name and header
                        seq_name = filename + "T" + str(start) + "-" +\
                                   str(start+window)+ "T" +\
                                   "{:.2f}".format(max_conf)
                        all_sequences.append(">"+seq_name+"\n"+high_conf_seq)
                        # set the name and header of the reversed sequence
                        seq_name_rev = seq_name + "Trev"
                        all_sequences.append(">"+seq_name_rev+"\n"+\
                                             high_conf_seq[::-1])

            # if the number of confidences is less than the window size,
            # just take the entire sequence with its confidences (on
            # condition that the average confidence of the sequence is
            # higher than the given threshold), and fill it left and right
            # with Xs so that the length of the sequence equals the size of
            # the window
            elif(np.average(conf) >= threshold):
                # get the number of required Xs left and right of the
                # sequence
                rest_x = (window - len(seq)) / 2
                # get the entire sequence filled with Xs left and right so
                # that the length of the sequence equals the window size
                # the left sided is rounded up, the right side rounded down
                high_conf_seq = math.ceil(rest_x)*"X" + "".join(seq) +\
                                math.floor(rest_x)*"X"
                # set the file name and header
                seq_name = filename + "T" + str(0) + "-" +\
                           str(len(high_conf_seq))+ "T" +\
                           "{:.2f}".format(np.average(conf))
                all_sequences.append(">"+seq_name+"\n"+ high_conf_seq)
                # set the name and header of the reversed sequence
                seq_name_rev = seq_name + "Trev"
                all_sequences.append(">"+seq_name_rev+"\n"+high_conf_seq[::-1])

    with open(os.path.join(output, "all_{}_collagens.mfaa".format(threshold)),
              "w") as writer:
              writer.write("\n".join(all_sequences))


if __name__ == "__main__":
    # path to the collagen modification file
    # set all non avaiable cells to -1
    mod_frame = pd.read_csv(sys.argv[4], sep=",", index_col=0)
    mod_frame = mod_frame.fillna(-1)
    # save the position of modifications
    # without_signal means that the signal sequence is removed from the
    # alpha-chain sequence if avaiable
    # only_chain means that only the chain sequence is taken from the alpha-chain
    # sequence
    # else, the entire sequence is taken
    mod_type = sys.argv[5]
    col_mods = None
    if(mod_type == "without_signal"):
        col_mods = {index:{"start":int(row["Signal"].split("-")[1]),
                           "end":row["Length"]} if row["Signal"] != -1
                           else {"start":0, "end":row["Length"]}
                    for index,row in mod_frame.iterrows()}
    elif(mod_type == "only_chain"):
        col_mods = {index:{"start":int(row["Chain"].split("-")[0])-1,
                           "end":int(row["Chain"].split("-")[1])}
                           for index,row in mod_frame.iterrows()}
    elif(mod_type == "whole_sequence"):
        col_mods = {index:{"start":0, "end":row["Length"]}
                    for index,row in mod_frame.iterrows()}
    else:
        raise Exception("Specify modification type")

    # user specifies the folder of the PDB files, length of the each
    # subsequence, a confidence threshold and the name of the output folder
    main(sys.argv[1], int(sys.argv[2]), int(sys.argv[3]), col_mods, sys.argv[6])
```
